# Massive outsourcing of energetically costly amino acids at the origin of animals

**DOI:** 10.1101/2024.04.18.590100

**Authors:** Niko Kasalo, Mirjana Domazet-Lošo, Tomislav Domazet-Lošo

## Abstract

Animals are generally capable of synthesizing eleven amino acids, while the remaining nine, often referred to as essential, must be acquired through diet. This characteristic profoundly impacts animals by defining their ecological lifestyles and evolutionary trajectory. Recent phylogenomic studies reveal that this phenotype results from gene losses that occurred at the root of the animal tree. However, it remains unclear which selective forces, if any, directed this far-reaching metabolic simplification event. Here, we show that essential amino acids are energetically far more expensive to synthesize than non-essential ones, particularly under high respiratory conditions—a hallmark of animal lifestyle. By applying permutation tests, we found that these difference in energy costs, counteracted by pleiotropy, created a selective pressure which led to the outsourcing of essential amino acids. Remarkably, we also found that extant animals use expensive amino acids more frequently compared to their closest unicellular relatives. This shows that animals significantly removed constraints on the usage of essential amino acids under high respiratory conditions by externalizing their production. Together, this implies that stabilizing selection underpinned by energy management drove this metabolic outsourcing in nutrient rich environments, thereby allowing animal genes to evolve more freely through protein sequence space. In this context, we propose that the origin of animals is tightly linked to energy-related adaptations rather than to unpredictable stochastic events, as recently suggested.

## Introduction

Proteins are made of 20 proteinogenic amino acids (AAs). The biosynthesis of these AAs is an indispensable biochemical process on which all life depends. Many organisms, such as plants, fungi, various bacteria, and many unicellular eukaryotes are capable of synthesizing all 20 AAs (Guedes et al. 2011; D’Souza et al. 2014; Trolle et al. 2022; Kanehisa et al. 2023). However, other lineages lost the capability to synthesize some AAs, which means that they have to harvest these essential amino acids (EAAs) from their environment (Guedes et al. 2011; D’Souza et al. 2014). A prominent example are animals (Metazoa), which lost the capability to synthesize nine EAAs (Guedes et al. 2011; Trolle et al. 2022; Payne & Loomis 2006). Recent phylogenomic studies showed that this simplification occurred at the root of the animal tree, after the split from choanoflagellates (Richter et al. 2018; Domazet-Lošo et al. 2024). This metabolic loss is part of a broader simplification event that preceded metazoan radiation and included a wide range of metabolic reductions (Richter et al. 2018; Domazet-Lošo et al. 2024; Guijarro-Clarke et al. 2020).

However, which selective forces, or random processes, were driving this simultaneous loss of EAA biosynthetic pathways in metazoans remains a great mystery (Trolle et al. 2022). The biosynthesis of AAs is generally energetically costly (Akashi & Gojobori 2002), but there is also considerable variability among them in the energy required for their production, with the cheapest and most expensive AAs differing by almost an order of magnitude (Akashi & Gojobori 2002; Wagner 2005; Zhang et al. 2018; Kaleta et al. 2013). On the other hand, free energy is a critical and limited resource that sustains biological systems, meaning that its harnessing and efficient usage is under continuous selective pressure (Judson 2017; Kepp 2020). For instance, energetically expensive AAs sparingly occur in protein sequences, especially in highly expressed genes (Akashi & Gojobori 2002; Seligmann 2003; Swire 2007; Heizer et al. 2011). It has been shown that this selectively favored bias leads to a net reduction in energy costs related to AA synthesis in cells (Judson 2017; Kepp 2020).

Inspired by these findings, we reasoned that the massive loss of EAA biosynthesis capability at the root of the animal tree could also be connected to energy management. A plausible hypothesis is that the biosynthesis of EAAs, as a group, incurs higher energy costs than the biosynthesis of non-essential AAs (NEAAs). By outsourcing the production of energetically costly AAs, an animal ancestor could have freed part of its energy budget, which could then be reallocated to other functional needs. We further assumed that the outsourcing of expensive EAAs would enable animals to use them more freely in their proteomes, compared to their unicellular holozoan ancestors, as the energetic constraints on their usage would be reduced. Surprisingly, these possibilities have never been tested, even though estimates of AA biosynthesis energy costs have been available for a long time (Akashi & Gojobori 2002; Wagner 2005; Zhang et al. 2018) and AA biosynthetic capabilities have been studied from various perspectives (Guedes et al. 2011; D’Souza et al. 2014; Trolle et al. 2022; Kanehisa et al. 2023; Payne & Loomis 2006; Zhang et al. 2018).

It is also puzzling that no effort has yet been made to conceptually link AA synthesis costs with their essentiality status, nor has any study considered the role of selection in shaping AA auxotrophies in animals. Here, we specifically focus on the selective forces and ecophysiological conditions that led to the essential/non-essential AA dichotomy. Our analyses reveal that energy management and pleiotropic effects exert antagonistic selective pressures, which — together with ecophysiological factors such as respiration mode and AA availability in the environment — led to the outsourcing of essential AAs in the metazoan lineage.

## Results

### Essential amino acids are expensive

We first retrieved the most recent estimates (Kaleta et al. 2013) on the required number of energy-bearing metabolites (ATP, NADH, and NADPH) for AA biosynthesis (Supplementary Data 1). These new estimates corrected some inadvertent errors that had propagated in previous studies using these values to calculate AA biosynthesis energy costs (see Methods). With these corrected values, we then calculated the direct cost of AA biosynthesis, representing the number of high-energy phosphate bonds (∼P) required for synthesizing an AA from its metabolic precursor (Supplementary Data 1, Supplementary Table 1).

It is important to note that direct costs quantify the energy used for AA production from metabolic precursors without considering how this metabolic decision impacts the overall energy balance of the cell. To address this, we also calculated the opportunity cost (see Methods), defined as the total energy that would have been produced if the precursors had been metabolized, plus the energy lost during the synthesis of AAs (Supplementary Data 1, Supplementary Table 1). This alternative cost measure, which has been extensively used in previous work (Craig & Weber 1998; Wagner 2005; Zhang et al. 2018; Akashi & Gojobori 2002), quantifies the impact of the decision to synthesize an AA on the overall energy balance of the cell by accounting for the unrealized energy gain from precursors (lost opportunity). In essence, it shows how much energy is given up by an organism when producing AAs (Craig & Weber 1998).

However, actual values of both opportunity costs and direct costs are directly dependent on the respiratory mode of a cell which can differ within the life cycle of an organism as well as between different species (Wagner 2005, Basan et al. 2015, Ferguson 2010, de Kok 2012). Surprisingly, the fundamental importance of respiration mode for AA energy cost estimation is rarely recognized (Wagner 2005, Kaleta et al. 2013). To account for this effect, we calculated here both direct and opportunity costs for three respiratory modes: fermentation, low respiration, and high respiration (see Methods, Supplementary Data 1, Supplementary Table 1). This range of respiration modes encompasses all metabolic situations across the tree of life, enabling us to identify the respiratory conditions that most favor the outsourcing of AAs.

For comparative purposes, we first sorted the 20 AAs in increasing order based on the opportunity cost calculated for high respiratory conditions, which correspond to the metabolic lifestyle of Metazoans (Fig. 1A). To illustrate how the dispensability status of AAs in animals is distributed in this representation, we marked the 9 EAAs and 11 NEAAs (Guedes et al. 2011; Trolle et al. 2022; Payne & Loomis 2006) (Fig. 1A). Remarkably, the AAs are arranged almost perfectly into two groups: (1) energetically cheaper NEAAs and (2) energetically more expensive EAAs (Fig. 1A).

**Fig. 1.**
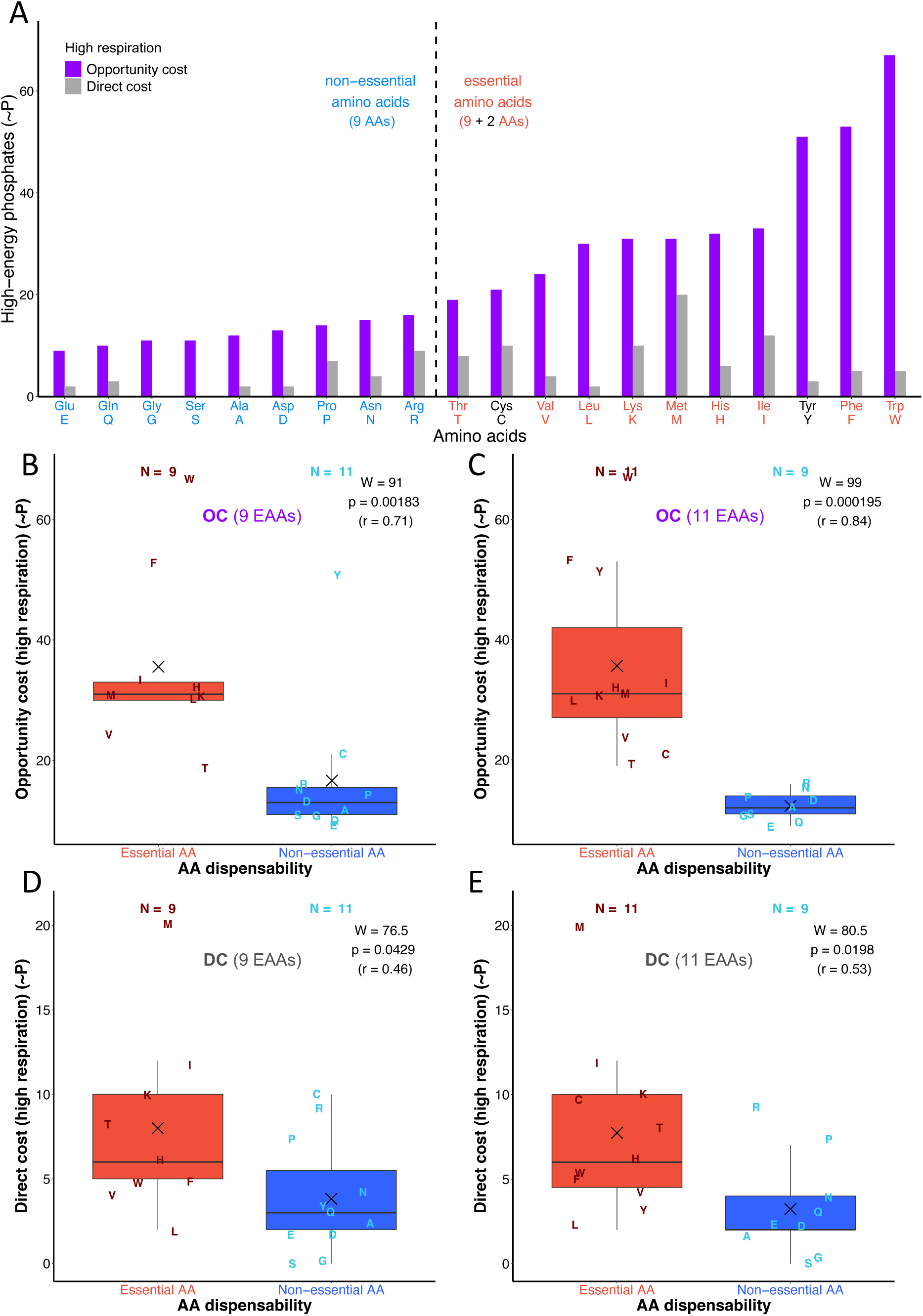
The comparison of amino acid synthesis costs. (**A)** The opportunity cost (OC) and direct cost (DC) of amino acid synthesis are shown as the number of high-energy phosphate bonds (∼P) required for AA synthesis. Amino acids are sorted in ascending order according to opportunity costs. (**B***–***E)** The comparison of energy costs between essential and non-essential amino acids in Metazoa. The opportunity cost represents the energy expended on AA synthesis combined with the energy that could be produced from precursor molecules, while the direct cost represents the energy expended on AA synthesis only (see Supplementary Table 1). We performed the comparison by considering 9 essential and 11 non-essential amino acids (**B, D**), and by considering 11 essential and 9 non-essential amino acids (**C, E**). In the extended AA dispensability analyses (**C, E**), we considered cysteine (C) and tyrosine (Y) as conditionally essential amino acids given that their synthesis directly depends on the essential amino acids methionine (M) and phenylalanine (F), respectively. The differences in energy costs between the two groups were shown by boxplots and the significance of these differences was tested by the Mann-Whitney U test with continuity correction. We depicted the corresponding W-value, p-value, and effect size (r) in each panel. The X symbol represents the mean. Individual AAs are shown by one-letter symbols.

The only exceptions are cysteine (C) and tyrosine (Y), which have relatively high energy costs and are usually considered NEAAs. However, in animals they are synthesized directly from other EAAs which are secured from the environment; i.e., cysteine (C) from methionine (M) and tyrosine (Y) from phenylalanine (F) (Kanehisa et al. 2018). This suggests that cysteine (C) and tyrosine (Y) should be considered partially essential as most of their synthesis costs are externally covered. In this context, it is evident that nearly all energy costs related to the top 55% (11 out of 20) most expensive AAs are outsourced (Fig. 1A). Analyses of opportunity costs under low respiration and fermentation conditions revealed very similar patterns; however, with overall lower values and lower difference between the cheapest and most expensive AAs (Supplementary Figs 1 and 2).

Although opportunity cost is the prevailing energy cost measure in previous studies (Craig & Weber 1998, Wagner 2005, Zhang et al. 2018, Akashi & Gojobori 2002), we also considered direct cost here. In contrast to opportunity costs, which account for the alternative option of respiring AA precursors, direct costs strictly reflect the expenses of AA synthesis. This measure is therefore useful in situations where cells have a surplus energy supply, making alternative metabolic choices less relevant. A simple visual inspection of this chart revealed that the direct costs of NEAAs are typically lower than those of EAAs; however, the difference between the two groups, although evident, is not as clear-cut as it is for opportunity costs (Fig. 1A, Supplementary Figs 1A and 2A). Thus, to assess these differences more quantitatively, we statistically compared average direct costs as well as opportunity costs between NEAAs and EAAs using a non-parametric test (Fig. 1B*–*E, Supplementary Figs 2B–E and 3B–E). We considered AA dispensability in two versions; in the first classical one, we grouped standard 9 EAAs (Fig. 1B,D), whereas in the second one, we extended this set to 11 members by treating cysteine (C) and tyrosine (Y) as EAAs (Fig. 1C,E).

Regardless of which cost measure is considered, under high respiration conditions the average energy costs of EAAs are significantly higher than the average energy costs of NEAAs with mostly large effect sizes (r > 0.5) (Fig. 1B*–*E). Interestingly, although average EAA costs are always higher than average NEAA costs; this difference becomes less apparent and significant under low respiration conditions and even more so under a fermentation lifestyle (Supplementary Figs 1 and 2). To better present this trend, we plotted the sum of AA opportunity costs as well as direct costs across three respiratory conditions (Fig. 2). This chart revealed a clear pattern where the differences in total AA synthesis costs between NEAAs and EAAs are the lowest under fermentation, increase under low respiration, and are the highest under high respiration (Fig. 2). This pattern is obvious for both direct and opportunity costs but is much stronger for opportunity costs, showing that more efficient respiration disproportionally increases the energy costs of EAAs in contrast to NEAAs. If we assume that energy consumption related to AA synthesis influences fitness, then the observed differences between NEAAs and EAAs should be most visible to selection under the conditions of high respiration. Subsequently, this selective pressure could lead to the eventual loss of EAA biosynthesis pathways, as observed at the root of Metazoa.

**Fig. 2.**
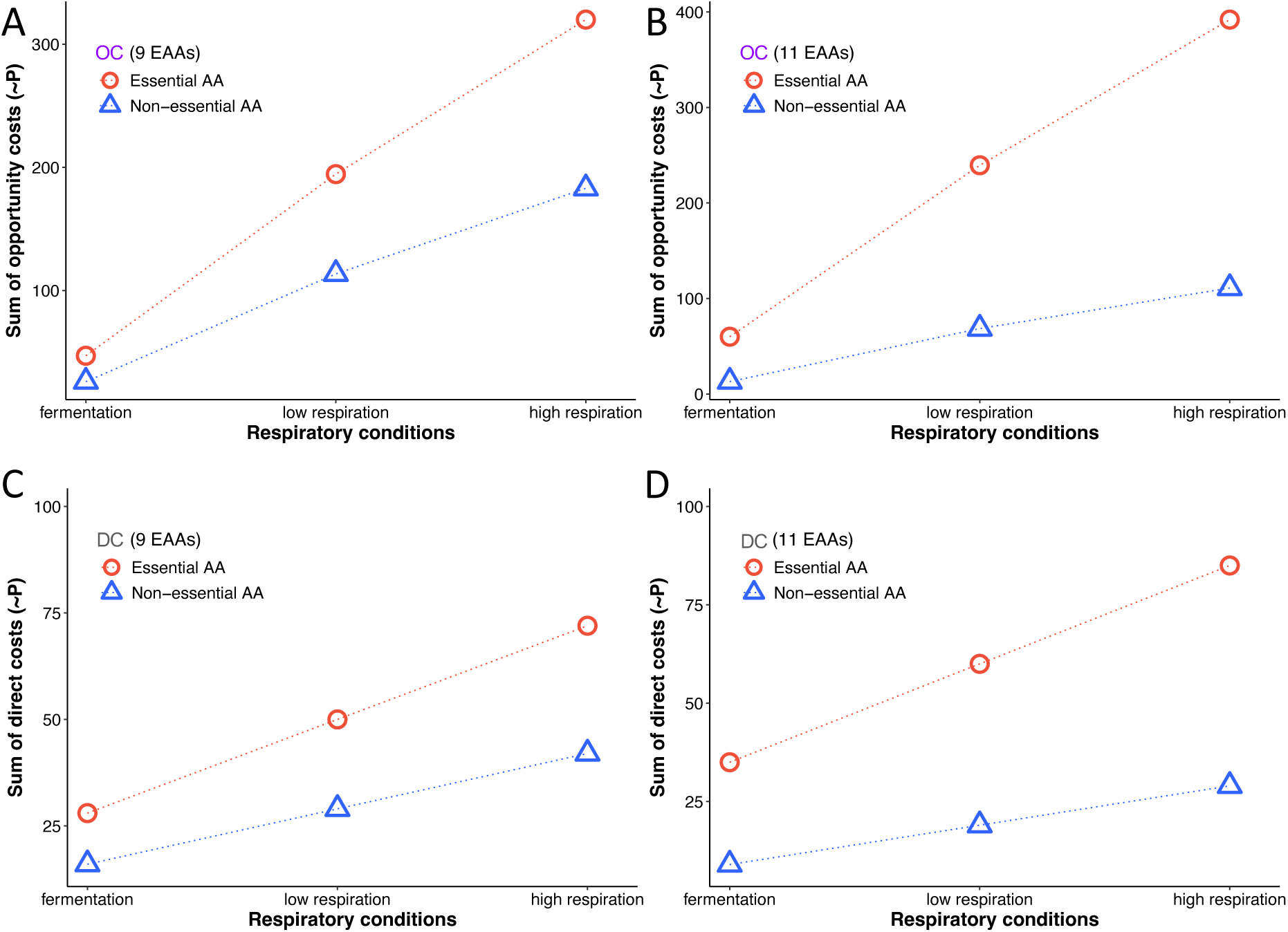
Dependence of amino acid biosynthesis costs on respiratory conditions. Summed opportunity costs (OC) and direct costs (DC) across all 20 AAs were calculated under fermentation, low respiration, and high respiration conditions (see Supplementary Table 1). Trends for EAAs (red line, circles) and NEAAs (blue line, triangles) are shown. The significance of differences between NEAAs and EAAs within a particular respiration mode are shown in Fig. 1, Supplementary Figs 1 and 2. We performed the calculations by considering 9 EAAs (**A, C**), and by considering 11 EAAs (**B, D**). In the extended AA dispensability analyses (**B, D**), we considered cysteine (C) and tyrosine (Y) as conditionally essential amino acids given that their synthesis depends on the essential amino acids methionine (M) and phenylalanine (F), respectively.

To further evaluate this hypothesis that selective forces related to energy management governed the outsourcing of EAAs under high respiration conditions, we devised a new test that could discriminate, in a probabilistic way, whether chance or selection was predominantly driving the evolution of the tested phenotype. We first generated all possible permutations of AA dispensability status under the assumption that 9 (or 11) out of 20 AAs are essential and then calculated the average opportunity (direct) cost of EAAs for every permutation. Based on the obtained distribution of average opportunity (direct) costs of EAAs, we calculated the matching empirical probability mass function (PMF) which was then used to retrieve the probability that the real-life set of EAAs in animals appeared randomly (Fig. 3). A low p-value would indicate that selection related to the AA synthesis costs shaped the observed set of EAAs in animals more strongly than random processes.

**Fig. 3.**
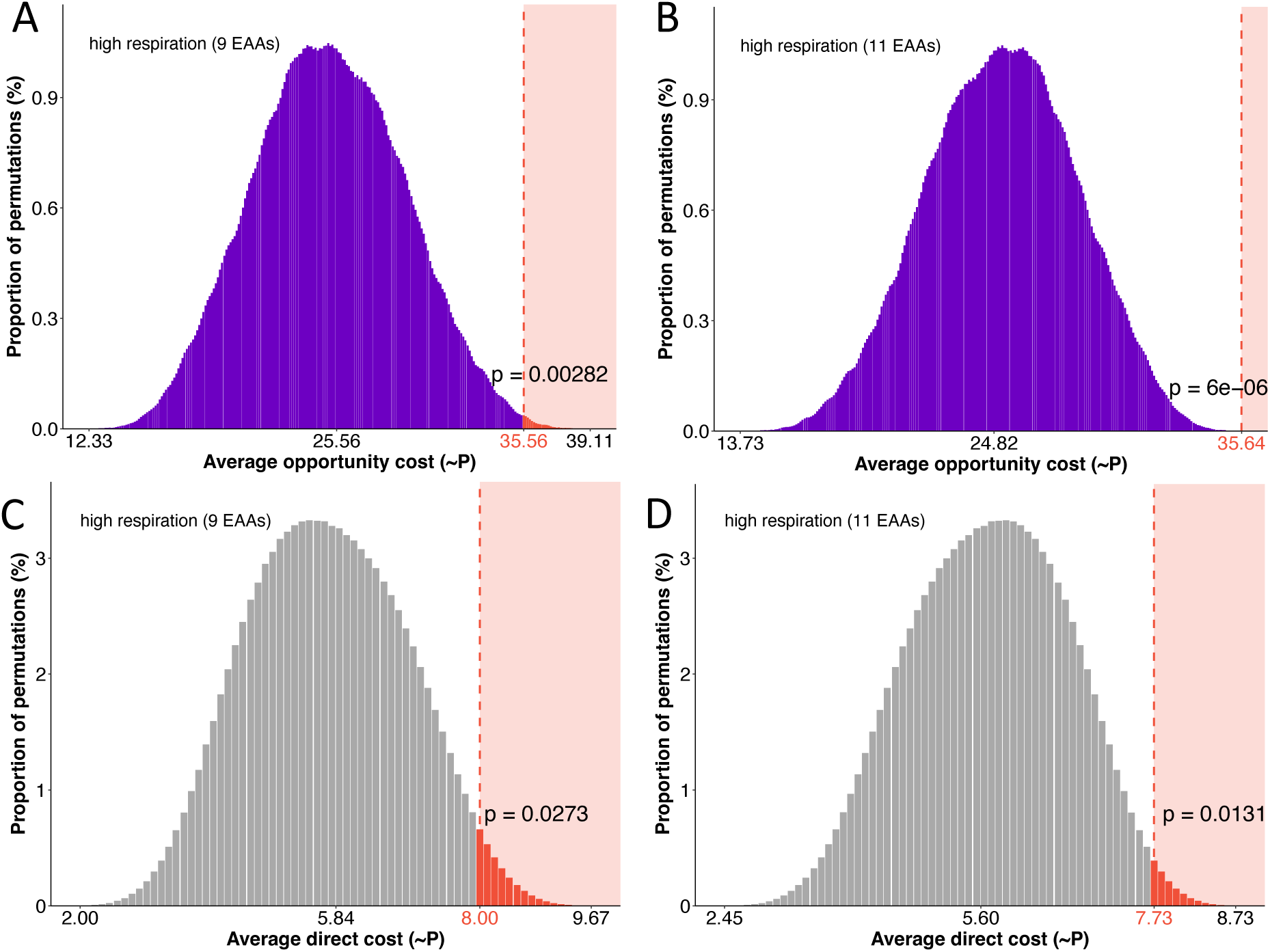
Permutation analyses of energy cost measures (selection tests). We assigned essentiality status to all possible combinations of 9 or 11 AAs (167,960 permutations). For each AA group in each permutation, average opportunity **(A, B)** and direct costs **(C, D)** were calculated, and the proportions of these averages are shown in histograms. The obtained distribution represents the empirical probability mass function (PMF). The value in red denotes the average value of the EAA sets observed in nature. The p-value was calculated by summing the proportions of average values equal to, or more extreme, than the observed average. Low p-values indicate high probability that selection pressures related to AA energy management draw the loss of EAAs synthesis capability.

All performed selection tests showed that the mean cost of the observed EAAs in animals falls at the right tail of the empirical distribution (Fig. 3). In fact, in some analyses, the observed EAA combination in animals has the highest average opportunity cost in comparisons to 167,960 possible permutations (Fig. 3B). This implies that it is extremely unlikely for such a combination of EAAs to arise by chance alone, and that AA production costs are an important selective force that shaped the evolution of the EAA set in animals. Although animal life-style is universally linked to high respiration, it is interesting to test if selection could shape the EAA set observed in animals under low respiration and fermentation conditions. Similar to high respiration data, permutation analysis of low respiration and fermentation data returned quite low p-values, which suggest that even in these metabolic situations, it is unlikely that random processes were a major force in shaping the EAA set. However, these p-values are less significant compared to the p-values obtained for cost measures under high respiration (Supplementary Fig. 3), which suggests that the selective loss of EAAs is the most probable under high respiratory conditions.

### Stabilizing selection via AA energetics and pleiotropy

If one assumes that AA energy management is an important selective force that draws the loss of EAAs production, the question arises as to why only half of the most expensive AAs were outsourced in animals. Theoretically, if AA energetics exert strong selective pressure, all AAs could potentially be outsourced. An expected answer to this question would be that AA pleiotropy counteracts AA energetics which together form favorable conditions for stabilizing selection. To test this idea, we applied the same battery of tests used in the analyses of AA energy costs. To visually inspect the data, we first mapped the number of KEGG pathways and biochemical reactions on individual AAs distributed according to their increasing opportunity costs (Fig. 4A). It can be seen that most of the cheapest AAs, and especially glutamate, take part in a large number of reactions and pathways. However, two conditionally essential AAs (cysteine and tyrosine) and half of the essential AAs also display relatively high involvement in various cellular processes (Fig. 4A).

**Fig. 4.**
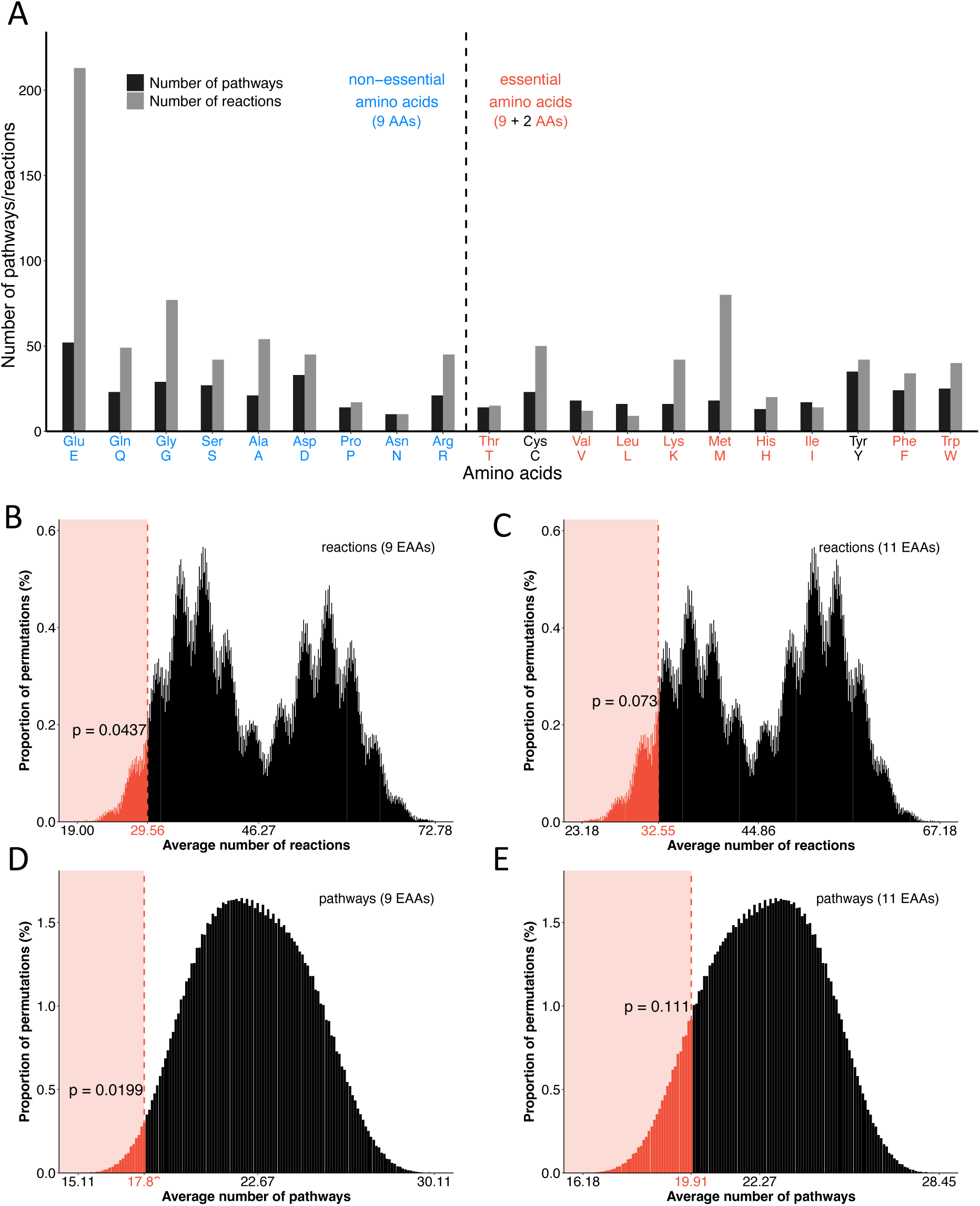
The pleiotropy measures and their permutation analyses (selection tests) (B*–*E). (**A)** For each amino acid, the number of pathways and reactions encoded in the KEGG database are shown. Amino acids are sorted in ascending order according to their opportunity costs. **(B*–* E)** We assigned essentiality status to all possible combinations of 9 or 11 AAs (167,960 permutations). For each AA group in each permutation, the average number of reactions **(B*–* C)** and pathways **(D*–*E)** were calculated and the proportions of these averages are shown in histograms (empirical PDF). The value in red denotes the average pleiotropy value of the EAA set observed in nature. The p-value was calculated by summing the proportions of average values equal to or lower than the observed average. Relatively low p-values indicate that selection pressures related to AA pleiotropy counteracted those related to AA energy management to some extent.

The comparison using non-parametric tests showed that EAAs have on average lower pleiotropy than NEAAs, however with borderline significance (Supplementary Fig. 4). To test if these differences are preferentially governed by random processes or selection, we again applied permutation analysis where the tested factor, instead of energy costs, was pleiotropy measured by the number of KEGG pathways and biochemical reactions in which AAs participate. This test showed that the averages of observed EAA pleiotropies fall at the left tails of empirical PMF with p-values that revolve on both sides of the 0.05 threshold (Fig. 4B-E). These results indicate that pleiotropy imposes selective pressure on maintaining the biosynthetic capabilities of some AAs. In turn, this also implies that pleiotropy counteracts energy-related selection to some extent. However, it seems that pleiotropy is a weaker force in this interaction.

### AA dispensability in animals

Nine AAs are canonically considered essential in animals, although there may be exceptions to this rule, often related to further losses in biosynthetic capabilities (Payne & Loomis 2006; Li et al. 2019; Lan 2021; de Oliveira 2022). However, the information on the AA dispensability status in animals and their allies is rather cursory and scattered over methodologically disparate studies (Lasser & Allen 1976; Cowey et al. 1970; Payne & Loomis 2006; Zečić et al. 2019; Li et al. 2019; Lan 2021; de Oliveira 2022; Richter et al. 2018). To get a coherent overview of AAs dispensability in animals and their closest relatives (non-animal holozoans), we explored the completeness of AA synthesis pathways on a large holozoan tree containing 167 species (Fig. 5, Supplementary Fig. 5). We note that the completeness of AA synthesis pathways is only a proxy of AA dispensability because it is possible that some taxa evolved or co-opted alternative pathways.

**Fig. 5.**
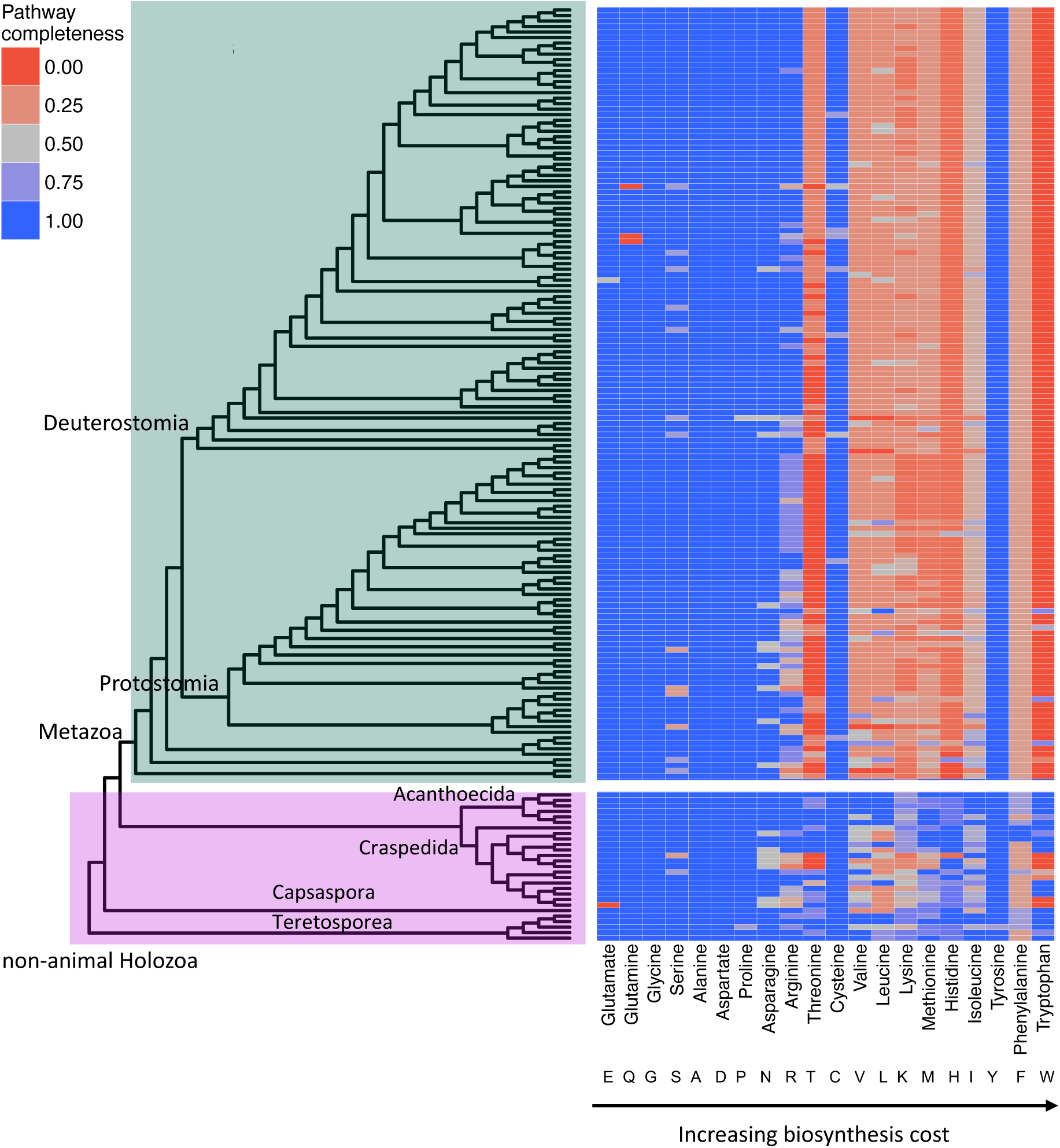
An overview of AA dispensability in Metazoa. We used the previously published database of 167 holozoans (Domazet-Lošo et al. 2024) to get a comprehensive overview of AA dispensability in this group. Fully resolved tree is shown Supplementary Fig. 5. We retrieved all enzymes involved in AA biosynthesis from the KEGG and MetaCyc databases. We searched for their homologs within our reference database using DIAMOND (see Methods). For each AA, we showed a completeness score, which represents the percentage of enzymes within a pathway that returned significant sequence similarity matches to our reference collection of AA biosynthesis enzymes. In the case of AAs with multiple alternative pathways, we showed the results only for the most complete one.

Most notably, we recovered an almost universal trend of reduction of pathways involved in the biosynthesis of the 9 canonically essential AAs in animals. This trend is not apparent in the non-animal holozoans where pathway completeness does not follow a clear pattern (Fig. 5, Supplementary Fig. 5). Since the choanoflagellate group Acanthoecida has in our analysis high completeness of biochemical pathways, and the same is true for Teretosporea and *Capsaspora*, we conclude that the ability to synthesize all 20 AAs is likely a plesiomorphic feature of holozoans, which agrees with previous work (Richter et al. 2018). In our analysis, we detected very few additional AA biosynthesis pathways with low completeness in animals beyond the canonical nine. The only exception is arginine, which is detected as partially complete in many non-vertebrate taxa (Fig. 5, Supplementary Fig. 5). This result aligns well with previous studies that report arginine auxotrophy in some insects and tunicates (Payne and Loomis 2006, Scaraffia et al. 2008). This volatility of auxotrophic/prototrophic status within animals is interesting since arginine is the most expensive NEAA; i.e., it is placed at border between NEAAs and EEAs when sorted by energy costs (Fig. 1A). Its dispensability status in animals is most likely dependent on the very sensitive interplay between taxon-specific differences in its environmental availability, the energy burden it poses on the metabolism and its pleiotropic effects (Payne and Loomis 2006, Scaraffia et al. 2008).

### Animal proteomes are expensive

The metabolic simplifications through the loss of EAA synthesis capabilities were only possible under the condition that the last common ancestor of animals (animal LCA) was able to secure in surplus these costly EAAs from the environment. If we assume that the unicellular ancestor of animals was a prototroph for AAs (Richter et al. 2018, Fig 5.), then some important shift in the feeding ecology occurred in the animal LCA that allowed this massive metabolic outsourcing. This might be a switch from a predominantly bacteria-eating lifestyle, as seen in present-day choanoflagellates (Ros-Rocher et al. 2021; Ruiz-Trillo et al. 2023) to more efficient multicellular-style suspension/filter feeding, or possibly to some sort of grazing behavior or macrophagous predation (Stanley 1973; Bengtson 2002).

Irrespective of which of these feeding adaptations occurred, it should have allowed animals to use EAAs more freely in their proteomes, since the energy for their production was largely outsourced. To test this idea, we compared the usage of EEAs between non-animal holozoans and Metazoa (Fig. 6, Supplementary Fig. 6). This comparison showed that animals use on average more EAAs in their proteins than non-animal holozoans (Fig. 6A and B). Similarly, we also found that the energy cost of an average AA is higher in metazoan than non-animal holozoans proteomes (Fig. 6C and D). However, if all genes are considered, the observed trends could hinge on the internal evolutionary dynamics of a particular lineage. To account for this, we reduced the set to include only the conserved genes shared between animals and non-animal holozoans (see Methods). The patterns obtained for these conserved genes are congruent with those obtained for all genes (Supplementary Fig. 6), suggesting that in animals, the restriction on the usage of expensive AAs is also relaxed for conserved genes. These results are in line with our prediction that the reduction of selective constraints on energetically expensive AAs allowed animals to more freely explore protein sequence space (Domazet-Lošo & Tautz 2003; Domazet-Lošo et al. 2007; Tautz & Domazet-Lošo 2011).

**Fig. 6.**
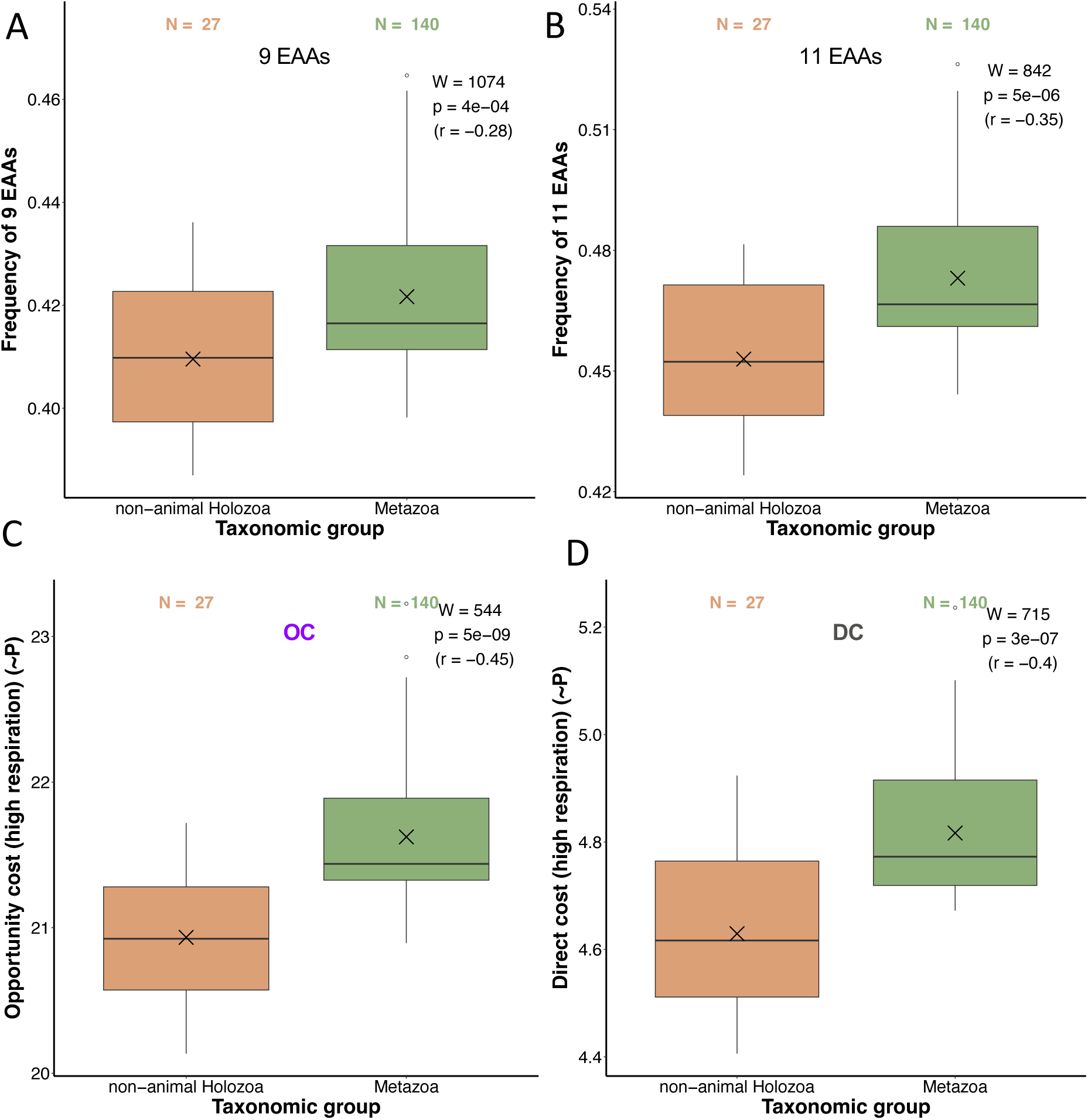
The comparison of EAA usage and the energy costs of an average AA in non-animal Holoza and Metazoa. (**A**) The cumulative frequency of nine AAs which are generally considered to be essential in Metazoans (Thr, Val, Leu, Lys, Ile, Met, His, Phe, Trp). (**B**) The cumulative frequency of eleven EAAs. This extended dataset contains two AAs which could be considered conditionally essential (Cys, Tyr). (**C***–***D**) The opportunity cost (OC) and the direct cost (DC) of an average AA (see Supplementary Table 1). This value represents a weighted mean of AA biosynthesis energy costs where the frequencies of twenty AAs in a proteome act as weights. The differences in energy costs between the two groups were shown by boxplots and the significance of these differences was tested by the Mann-Whitney U test with continuity correction. We depicted the corresponding W-value, p-value, and effect size (r) in each panel. The X symbol represents the mean. The list of non-animal holozoans (27) and metazoans (140) whose proteomes are included in calculations is available in Supplementary Data 2 (Domazet-Lošo et al. 2024).

## Discussion

Several studies examined the genes involved in the AA biosynthesis pathways and found that many of them are missing in animals (e.g., Guedes et al. 2011; Payne & Loomis 2006; de Oliveira et al. 2022, Richter et al. 2018; Domazet-Lošo et al. 2024). It was suggested that these reductions are caused by animal heterotrophy that relaxed selective pressures on maintaining AA biosynthetic capabilities (Payne & Loomis 2006, Guedes et al. 2011). While improved heterotrophic feeding strategies likely played a role in the outsourcing of EAAs in animals, this is just one factor in the presumably more complex scenario that resulted in this metabolic simplification. For instance, it has not been investigated which evolutionary forces primarily drove the loss of amino acid synthesis capabilities after purifying selection for their maintenance was reduced due to improved dietary access to amino acids. Some studies vaguely suggest that random processes contributed to the loss of dispensable amino acid synthesis pathways (Payne & Loomis 2006, Guedes et al. 2011); however, the possibility of selectively driven EAA loss has not been considered. To address this, we propose an amino acid outsourcing model based on the findings of this study (Fig. 7).

**Fig. 7.**
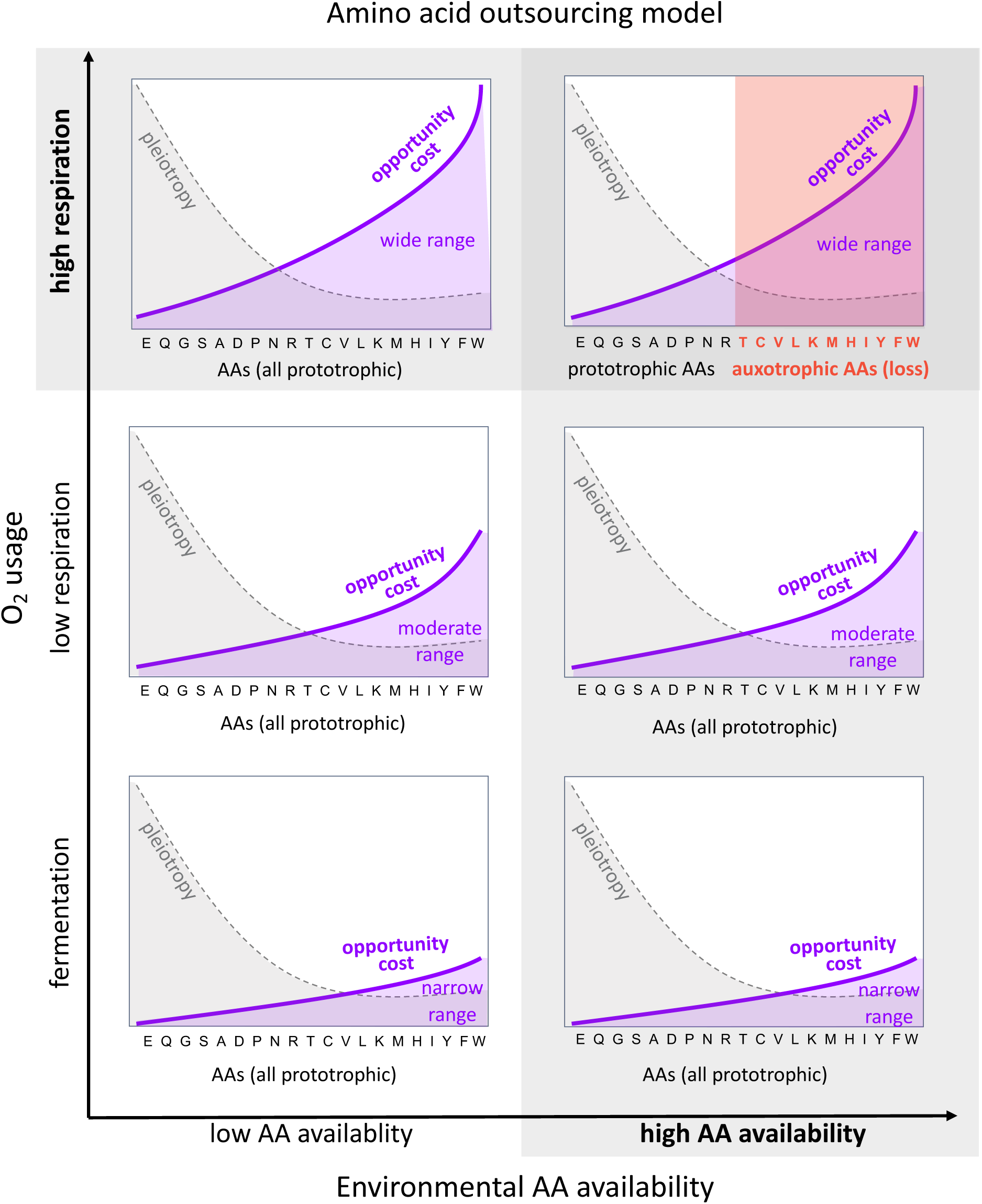
Amino acid outsourcing model. Six charts depict the interplay between key predictors of AA auxotrophy: (i) AA biosynthesis costs, (ii) pleiotropy, (iii) the availability of AAs in the environment, and (iv) respiration mode. On each chart, AAs are ordered by increasing opportunity costs (OC), which are depicted by the purple line. The area below the purple line depicts the total AA opportunity costs. The degree of change between the lowest and highest OC value is designated as ‘narrow’, ‘moderate’ and ‘high range’. The dashed line depicts the pleiotropic effects of AAs, with higher values indicating more reactions and pathways in which the AA takes part. Shaded backgrounds indicate ecophysiological conditions that favor the loss of AA prototrophy. All graphs are simplified drawings of data presented in this study.

According to our model, the availability of environmental amino acids and an organism’s respiration mode are primary ecophysiological determinants that set the stage for the outsourcing of AAs (Fig. 7). A surplus of amino acids in the environment reduces selective pressures on in-house amino acid production (Fig. 7) by lowering the net energy costs associated with this production. However, the extent of these savings largely depends on the respiration physiology of an organism that resides in an AA-rich environment (Fig. 7). The total costs of AA production substantially increase with the shift from fermentation to high respiration (Fig. 2, Supplementary Table 1). This increase applies to both direct costs and opportunity costs, with the total for all 20 AAs rising by 2.6 times and 6.9 times, respectively, as the respiration mode shifts from fermentation to high respiration (Fig. 2, Supplementary Table 1). This dramatic increase indicates that selective pressures related to AA production costs are substantially higher in organisms, such as animals, that have adopted a high-respiration lifestyle. Additionally, the metabolic shift from fermentation to high respiration widens the differences between the cheapest and most expensive AAs (Fig. 2, Fig. 7).

All of this suggests that the strongest energy-related selective pressures will act on organisms thriving in oxygenated environments where AAs are readily available and whose ecophysiology relies on high respiration. Beyond animals, at least two other examples of convergence in eukaryotic lineages support this model. The soil-dwelling amoeba *Dictyostelium discoideum* (11 EAAs) and the euglenozoan *Leishmania major* (also 11 EAAs) demonstrate a striking similarity to metazoans regarding amino acid dispensability, with only serine and arginine being additionally essential (Payne & Loomis 2006). Both organisms thrive in oxygen-rich environments (Cochet-Escartin et al. 2021, Duarte et al. 2021), and their feeding ecologies — *L. major* as a parasite and *D. discoideum* as a generalist predator (Shreenidhi et al. 2024) — resemble those of certain animal lineages. These ecological parallelisms help to explain this convergence in AA auxotrophy. Furthermore, it was experimentally demonstrated that the bacterial strains that are auxotrophic for various amino acids gain a fitness advantage over their prototrophic counterparts in AA-rich cultures (D’Souza et al. 2014), indicating that selective pressure on energy management universally underpins AA auxotrophies.

On the other hand, fully AA-prototrophic *E. coli* strains utilize an ‘overflow’ mechanism to genetically switch from aerobic to anaerobic metabolism (Basan et al., 2015). This approach enables cheaper AA production during periods of high cell proliferation and protein synthesis, when demand is high. This highlights that, while an aerobic lifestyle generally allows for more efficient energy production, it also incurs the cost of more expensive AA synthesis.

There are at least three solutions to this persistent trade-off between the overall energy budget and AA production under aerobic conditions. One strategy is to outsource the production of energetically expensive AAs, as seen in all animals, as well as in *D. discoideum* and *L. major*. Another solution is the ‘overflow’ mechanism based on gene regulation, as observed in *E. coli*. The third approach is a shift to a predominantly fermentative lifestyle with occasional aerobic episodes featuring reduced respiration, as seen in some yeasts (Marcet-Houben et al., 2009; Domazet-Lošo et al., 2024).

In line with this, our permutation analyses, which probabilistically distinguish the relative contributions of selection and random processes, strongly suggest that selective pressure related to cost optimization primarily drove the loss of capability for EAA production (Fig. 3, Fig. 7). However, further permutation-based tests also indicated that this energy-driven selection was somewhat counteracted by pleiotropy as a selective force, which prevented the loss of synthesis capabilities for NEAAs that have important pleiotropic functions (Fig. 4, Fig. 7). For instance, glutamate (E), the most pleiotropic amino acid, serves as a major metabolic hub, playing a critical role in nitrogen assimilation, nucleoside and amino acid biosynthesis (Walker & van der Donk 2016). Related glutamine (Q), which is synthesized by glutamine synthetase from glutamate and ammonia, also has diverse metabolic functions, including nitrogen metabolism, nucleotide synthesis, non-essential amino acid synthesis, and the regulation of epigenetic changes (Yoo et al. 2020). Interestingly, glutamine synthetase is one of the oldest known functional genes, present in the last universal common ancestor, suggesting its fundamental role in cellular metabolism (Carvalho Fernandes et al. 2022). Comparably to glutamate (E) and glutamine (Q), other NEAAs are also known to perform a variety of important metabolic functions besides protein synthesis (Krall et al. 2016, Zhu et al. 2017, Holeček 2022, Christgen & Becker 2019, Scaraffia et al. 2008).

It should also be emphasized that animals still utilize all AAs. However, energetically expensive ones (EAAs) are sourced externally, allowing animals to conserve the energy that would be required for their production. We previously introduced the term “functional outsourcing” to describe this phenomenon, since simplified organisms still rely on the lost biosynthetic pathways, which are now outsourced to the environment (Domazet-Lošo et al. 2024). This phenomenon is modeled by the Black Queen Hypothesis, which explains the evolution of ecological dependencies arising from gene loss. According to this hypothesis, some free-living organisms can lose a costly function if it is continuously provided by other organisms in the accessible environment (Morris et al. 2012, Mas et al. 2016).

Following this argumentation line, we think that the terms “essential” and “non-essential” AAs, which were coined within the context of animal dietary science, are quite misleading. All twenty proteinogenic amino acids are indispensable for every cellular organism on earth, and therefore essential. In fact, most of the “non-essential” AAs have to be supplemented to some extent by feeding (Wu et al. 2014). Therefore, the only relevant question is whether they are internally or externally synthesized, or in other words, built in-house or outsourced (Domazet-Lošo et al. 2024). We thus advocate that the terms auxotrophic amino acids (AAAs) and prototrophic amino acids (PAAs) should be adopted when the dispensability of AAs is discussed, regardless of which organism on the tree of life is in focus.

In summary, we have shown that the biosynthesis of nutritionally essential AAs is far more expensive than that of non-essential ones, regardless of the cost measures applied. We also found strong support for the idea that the loss of EAA biosynthesis capability is an important synapomorphy of animals that is primarily driven by natural selection related to energy-saving requirements. Around 575 million years ago, Earth’s ecosystems were profoundly impacted by the oxygenation of oceans (Brocks et al. 2017) and new ecological interactions that emerged together with the rise of animals and their heterotrophic way of life (Judson 2017). We propose that stabilizing selection, driven by energy economy and balanced by pleiotropy, favored the loss of expensive AA biosynthesis until an equilibrium between functional outsourcing and functional autonomy was reached, resulting in the so-called essential and non-essential AA sets.

This important selection-driven metabolic outsourcing allowed animals to more freely use energetically expensive AAs in already existing and newly arisen genes (Domazet-Lošo et al. 2024; Domazet-Lošo & Tautz 2003; Domazet-Lošo et al. 2007; Tautz & Domazet-Lošo 2011). Whether this opportunity also allowed animals to evolve functionally more optimized proteins has yet to be explored. We also suspect that the first animals, by outsourcing the production of expensive amino acids, opened a window of opportunity which allowed them to relocate a substantial part of their energy budget from costly metabolic synthesis to more animal-specific and energy-demanding functions, such as muscle movements, electric signaling, and increased protein glycosylation (Kifer et al. 2024). In this context, we think that the recent suggestion that the origin of animals was a stochasticity-guided black swan event (Ruiz-Trillo et al. 2023) is probably premature.

## Methods

### AA cost estimates

We obtained data on the number of ATP and NAD(P)H used in the biosynthesis of AAs from Kaleta et al. (2013). Previous studies primarily relied on estimates from Craig & Weber (1998), which contain some inaccuracies. For instance, ATP values for asparagine and serine were erroneously transcribed from the original source (Neidhardt et al. 1990). Additionally, minor updates were made to the ATP and NAD(P)H values for arginine, cysteine, histidine, methionine, and tryptophan based on the most recent biochemical findings (Kaleta et al. 2013). We also retrieved the number of ATP and NAD(P)H utilized in the metabolism of AA precursors (Voet et al. 2016). To approximate ATP generation under different respiratory conditions, we converted the reducing equivalents to ATP as follows: (i) ‘high respiration,’ representing fully functional oxidative phosphorylation: 1 NAD(P)H = 2 FADH2 = 2 ATP (Kaleta et al. 2013); (ii) ‘low respiration,’ representing oxidative phosphorylation without proton pumping at complex I: 1 NAD(P)H = 2 FADH2 = 1 ATP, which corresponds, for instance, to the metabolism of *S. cerevisiae* and some *E. coli* strains (Schuetz et al. 2007; de Kok et al. 2012); (iii) ‘fermentation,’ representing anaerobic conditions without conversion of reducing equivalents to ATP.

We calculated the costs of AA production in two ways: direct cost and opportunity cost. The direct cost was calculated by summing ATP and ATP equivalents of energy-bearing metabolites (NADH and NADPH) used in the production of an AA, starting from the precursor molecules (Supplementary Table 1, Supplementary Data 1). The opportunity cost was calculated by summing the energy lost in the synthesis of AAs (direct cost) and the energy that would have been produced if a cell catabolizes precursors instead of making AAs (Supplementary Table 1, Supplementary Data 1). The energy gained from the catabolism of precursors was calculated by summing produced ATP and ATP equivalents of energy-bearing metabolites (NADH, NADPH, and FADH2) (Supplementary Table 1, Supplementary Data 1). The opportunity cost measure, which has been extensively used in previous work (Craig & Weber 1998, Wagner 2005, Zhang et al. 2018, Akashi & Gojobori 2002), quantifies how a decision to synthesize an AA impacts the overall energetic balance of a cell by taking into account an unrealized energy gain from precursors (lost opportunity). In essence, opportunity cost shows how much energy is given up by an organism when producing AAs (Craig & Weber 1998), and is analogous to the ‘opportunity cost’ concept in economics (Mankiw 2019).

### Bioinformatics and statistical analyses

The proteomes of non-animal holozoans (27 species) and metazoans (140 species) were taken from our previous study (Supplementary Data 2; Domazet-Lošo et al. 2024). Using this dataset, we calculated the frequency of each of the 20 AAs in each proteome. By summing these frequencies, we calculated the cumulative frequency of nine AAs which are generally considered to be essential in Metazoa (Thr, Val, Leu, Lys, Ile, Met, His, Phe, Trp). We repeated this analysis by adding two additional AAs that could be considered conditionally essential (Cys, Tyr). Using AAs frequencies, we also calculated the direct cost and opportunity cost of an average AA in each proteome. This value represents a weighted mean of AA biosynthesis costs where the frequencies of twenty AAs in a proteome are acting as weights (Supplementary Data 2).

To control for the presence of lineage-specific genes, we repeated this procedure using only genes conserved between animals and non-animal holozoans. We first clustered all 167 proteomes using the following MMSeqs2 parameters: -e 0.001 -c 0.8 --max-seqs 400 --cluster-mode 1 --cov-mode 0. By using a cutoff of c = 0.8, we obtained clusters of proteins with highly similar architectures (Domazet-Lošo et al. 2024). We then extracted only the clusters that contained both metazoan and non-metazoan holozoan proteins and repeated the frequency and energy cost analyses. The initial dataset contained 3,514,971 proteins, while the dataset with only conserved genes contained 1,460,551 proteins (42%).

Data on pleiotropy was collected from the KEGG database (Kanehisa et al., 2023) and is represented by two metrics: (i) the number of KEGG reactions, and (ii) the number of KEGG pathways in which each amino acid (AA) is involved. The significance of differences in the energy costs between EAAs and NEAAs, differences in the frequency of EAA usage, differences in the energy cost of an average AA, and differences in the number of reactions and pathways was tested by the two-tailed Mann-Whitney U with continuity correction. For each plot, we report the Mann-Whitney statistic (W-value), the p-value, and the effect size r.

The permutation analyses were performed by calculating the average of each measure (direct cost, opportunity cost, number of pathways, number of reactions) for each of the 167,960 possible permutations of 9 EAAs or 11 EAAs. Since there is a limited number of possible average values, we arranged the averages into bins containing identical values. To quantify this, we calculated the proportion of permutations resulting in that value by dividing the number of elements in the bin by the total number of permutations. The obtained distribution represents empirical probability mass function (PMF) which was then used to retrieve the probability that the real-life set of EAAs in animals appeared randomly. We calculated these propabilities (p-values) by summing the proportions of permutation in the range from the actual value observed in nature to the most extreme value at the closest distribution tail.

To construct the heatmap depicting the completeness of AA biosynthetic pathways, we first assembled a list of all enzymes involved in these pathways from the MetaCyc (Caspi et al. 2019) and KEGG (Kanehisa et al. 2023) databases. For AAs that can be synthesized by multiple alternative pathways, we annotated each pathway separately even if they share some enzymes.

In the next step, we compiled a list of organisms whose annotated genomes are available in the KEGG database (Supplementary Data 2). We selected organisms from all the major clades present in our database plus some additional eukaryotic organisms to increase the probability of finding a matching enzyme in non-animal holozoans (Supplementary Data 2). We then extracted the amino acid sequences of all enzymes involved in AA biosynthesis for each of these organisms, resulting in a total of 14797 sequences (Supplementary Data 2).

We used these sequences as queries to perform a DIAMOND protein sequence similarity search against our database of 167 holozoan proteomes (Domazet-Lošo et al. 2024). The following DIAMOND blastp parameters were applied: --ultra-sensitive --masking seg –evalue 0.001 --max-target-seqs 100000000 (Buchfink et al. 2021). To avoid false positives, we subsequently removed hits with a sequence identity of less than 40%. For each pathway and species in the database, we divided the number of enzymes detected by DIAMOND by the total number of enzymes present in that pathway, resulting in pathway completeness values ranging from 0 to 1. If alternative biosynthetic pathways were present in a species, we selected the most complete one.

The Mann-Whitney U tests were conducted in the R environment using the package rcompanion (https://CRAN.R-project.org/package=rcompanion). Permutations were calculated using the RcppAlgos package (https://CRAN.R-project.org/package=RcppAlgos). Phylogeny mapping was performed using the ggtree package (Yu et al. 2017).

## Data availability

All data are available in the main text or the supplementary materials.

## Author Contributions

T.D.-L. and M.D.-L. initiated the study. N.K., M.D.-L., T.D.-L. conceptualized and performed the analyses. N.K., prepared the figures and tables for publication. N.K., M.D.-L., T.D.-L. wrote the manuscript.

## Acknowledgments

We thank M. Futo, A. Tušar, S. Koska, D. Franjević and G. Klobučar for discussions. This work was supported by the Croatian Science Foundation under the project IP-2016-06-5924 (T.D.-L.), the City of Zagreb (T.D.-L.), the Adris Foundation (T.D.-L.), the European Regional Development Fund KK.01.1.1.01.0009 DATACROSS (M.D.-L., T.D.-L.). We used the computational resources of the University Computing Center - SRCE (Padobran) and the Institute Ruđer Bošković.

---

**Supplementary Table 1.** Amino acid biosynthesis costs calculated in two ways (direct and opportunity cost) for three different respiratory conditions (fermentation, low respiration and high respiration). Direct costs are estimated as the number of high-energy phosphate bonds (∼P) required for the synthesis of an AA from its metabolic precursors, while opportunity costs are estimated as the sum of direct costs and the energy that would have been gained from the precursors if they were metabolized. Canonical dietary essentiality of amino acids in animals is denoted by abbreviations NE (non-essential) and E (essential), while CE denotes two conditionally essential AAs (tyrosine and cysteine). Total denotes the sum of respective costs over all 20 AAs. The values in parentheses show fold change increase under respiratory conditions in comparison to fermentation. For more details on the computation of cost measures see Supplementary Data 1.

| AA name | Dispensability | Direct costs (~P) |  |  | Opportunity costs (~P) |  |  |
| --- | --- | --- | --- | --- | --- | --- | --- |
|  |  | fermen. | low resp. | high resp. | fermen. | low resp. | high resp. |
| Glutamate, E | NE | 0 | 1 | 2 | 0 | 5.5 | 9 |
| Glutamine, Q | NE | 1 | 2 | 3 | 1 | 6.5 | 10 |
| Glycine, G | NE | 0 | 0 | 0 | 1 | 6.5 | 11 |
| Serine, S | NE | 0 | 0 | 0 | 1 | 6.5 | 11 |
| Alanine, A | NE | 0 | 1 | 2 | 0 | 6.5 | 12 |
| Aspartate, D | NE | 0 | 1 | 2 | 1 | 7.5 | 13 |
| Proline, P | NE | 1 | 4 | 7 | 1 | 8.5 | 14 |
| Asparagine, N | NE | 2 | 3 | 4 | 3 | 9.5 | 15 |
| Arginine, R | NE | 5 | 7 | 9 | 5 | 11.5 | 16 |
| Threonine, T | E | 2 | 5 | 8 | 3 | 11.5 | 19 |
| Cysteine, C | NE, CE | 6 | 8 | 10 | 7 | 14.5 | 21 |
| Valine, V | E | 0 | 2 | 4 | 0 | 13 | 24 |
| Leucine, L | E | 0 | 1 | 2 | 0 | 16.5 | 30 |
| Lysine, K | E | 2 | 6 | 10 | 3 | 18 | 31 |
| Methionine, M | E | 10 | 15 | 20 | 11 | 21.5 | 31 |
| Histidine, H | E | 6 | 6 | 6 | 9 | 21.5 | 32 |
| Isoleucine, I | E | 2 | 7 | 12 | 3 | 19 | 33 |
| Tyrosine, Y | NE, CE | 1 | 2 | 3 | 6 | 30.5 | 51 |
| Phenylalanine, F | E | 1 | 3 | 5 | 6 | 31.5 | 53 |
| Tryptophan, W | E | 5 | 5 | 5 | 12 | 42 | 67 |
| Total |  | 44 | 79 (1.8) | 114 (2.6) | 73 | 308 (4.2) | 503 (6.9) |

**Supplementary Fig. 1.**
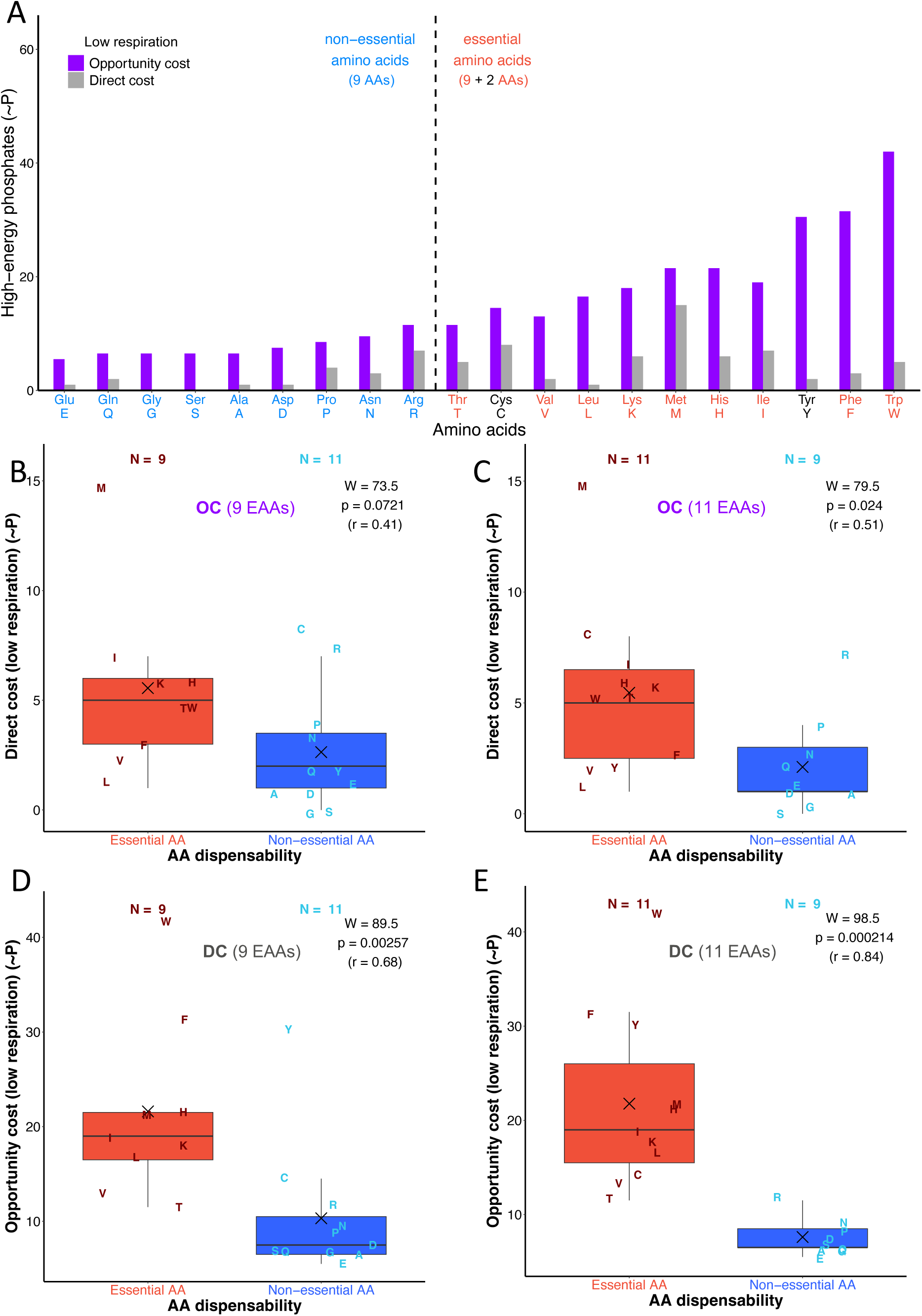
The comparison of amino acid synthesis costs under low respiratory conditions. (**A)** The opportunity cost (OC) and direct cost (DC) of an amino acid synthesis are shown as the number of high-energy phosphate bonds (∼P) required for AA synthesis. Amino acids are sorted in ascending order according to opportunity costs. (**B***–***E)** The comparison of energy costs between essential and non-essential amino acids in Metazoa. The opportunity cost represents the energy expended on AA synthesis combined with the energy that could be produced from precursor molecules, while the direct cost represents the energy expended on AA synthesis only (see Supplementary Table 1). We performed the comparison by considering 9 essential and 11 non-essential amino acids (**B, D**), and by considering 11 essential and 9 non-essential amino acids (**C, E**). In the extended AA dispensability analyses (**C, E**), we considered cysteine (C) and tyrosine (Y) as conditionally essential amino acids given that their synthesis directly depends on the essential amino acids methionine (M) and phenylalanine (F), respectively. The differences in energy costs between the two groups were shown by boxplots and the significance of these differences was tested by the Mann-Whitney U test with continuity correction. We depicted the corresponding W-value, p-value, and effect size (r) in each panel. The X symbol represents the mean. Individual AAs are shown by one-letter symbols.

**Supplementary Fig. 2.**
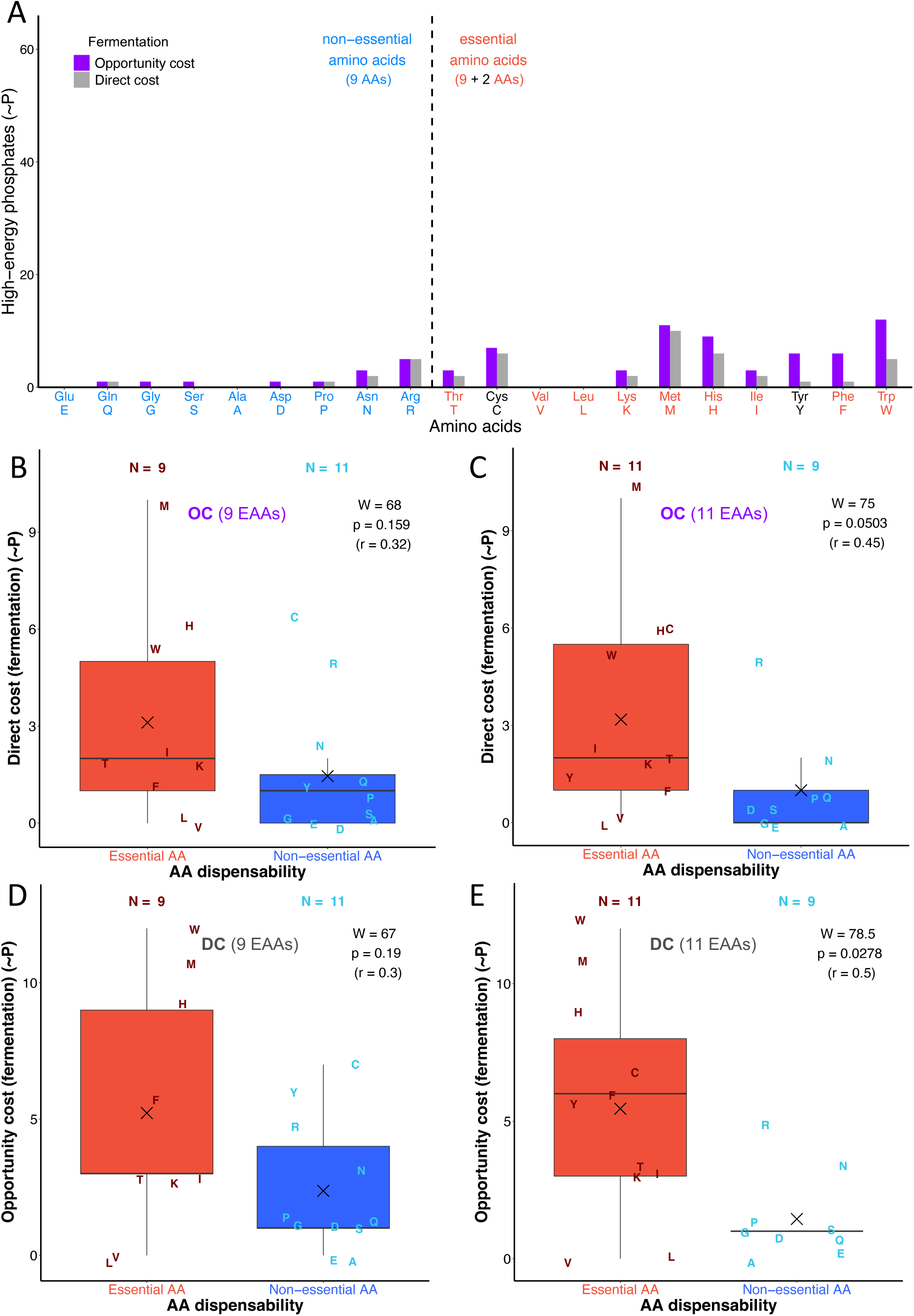
The comparison of amino acid synthesis costs under fermentative conditions. (**A)** The opportunity cost (OC) and direct cost (DC) of an amino acid synthesis are shown as the number of high-energy phosphate bonds (∼P) required for AA synthesis. Amino acids are sorted in ascending order according to opportunity costs. (**B***–***E)** The comparison of energy costs between essential and non-essential amino acids in Metazoa. The opportunity cost represents the energy expended on AA synthesis combined with the energy that could be produced from precursor molecules, while the direct cost represents the energy expended on AA synthesis only (see Supplementary Table 1). We performed the comparison by considering 9 essential and 11 non-essential amino acids (**B, D**), and by considering 11 essential and 9 non-essential amino acids (**C, E**). In the extended AA dispensability analyses (**C, E**), we considered cysteine (C) and tyrosine (Y) as conditionally essential amino acids given that their synthesis directly depends on the essential amino acids methionine (M) and phenylalanine (F), respectively. The differences in energy costs between the two groups were shown by boxplots and the significance of these differences was tested by the Mann-Whitney U test with continuity correction. We depicted the corresponding W-value, p-value, and effect size (r) in each panel. The X symbol represents the mean. Individual AAs are shown by one-letter symbols.

**Supplementary Fig. 3.**
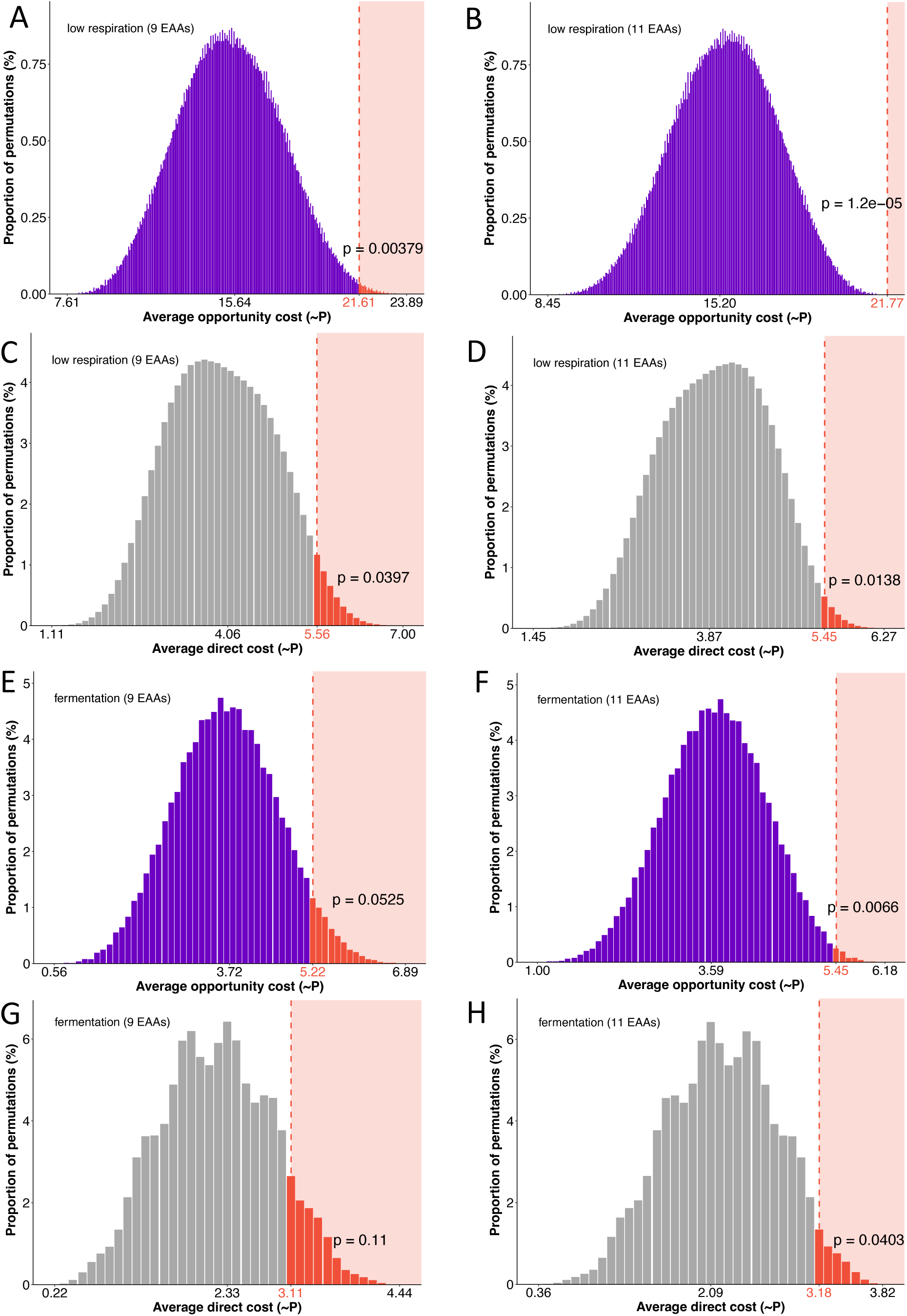
Permutation analyses of energy cost measures (selection tests) under low respiratory and fermentative conditions. We assigned essentiality status to all possible combinations of 9 or 11 AAs (167,960 permutations). For each AA group in each permutation, average opportunity costs **(A, B, E, F)** and direct costs **(C, D, G, H)** were calculated, and the proportions of these averages are shown in histograms. Energy cost under low respiration **(A-D)** and fermentation **(E-H)** were considered. The obtained distribution represents the empirical probability mass function (PMF). The value in red denotes the average value of the EAA sets observed in nature. The p-value was calculated by summing the proportions of average values equal to, or more extreme, than the observed average. Low p-values indicate high probability that selection pressures related to AA energy management draw the loss EAAs synthesis capability.

**Supplementary Fig. 4.**
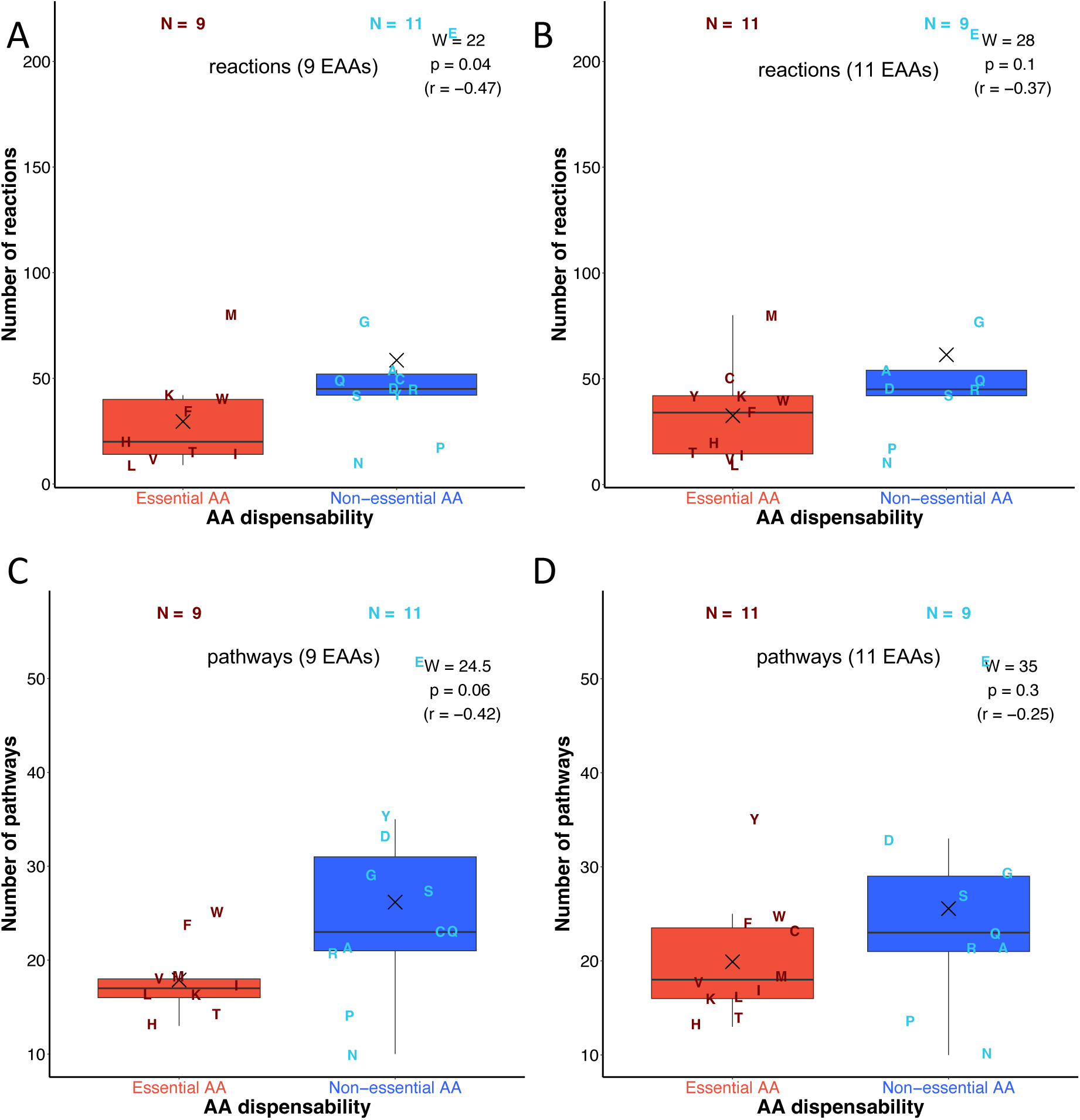
The comparison of amino acid pleiotropy. The comparison of the number of reactions and pathways between essential and non-essential amino acids in Metazoa. We performed the comparison by considering 9 essential and 11 non-essential amino acids (**A, C**), and by considering 11 essential and 9 non-essential amino acids (**C, E**). In the extended AA dispensability analyses (**B, D**), we considered cysteine (C) and tyrosine (Y) as conditionally essential amino acids given that their synthesis directly depends on the essential amino acids methionine (M) and phenylalanine (F), respectively. The differences in number of reactions and pathways between the two groups were shown by boxplots and the significance of these differences was tested by the Mann-Whitney U test with continuity correction. We depicted the corresponding W-value, p-value, and effect size (r) in each panel. The X symbol represents the mean. Individual AAs are shown by one-letter symbols. We acquired the number of metabolic pathways and reactions in which each AA is involved from the KEGG database (Kanehisa et al. 2023).

**Supplementary Fig. 5.**
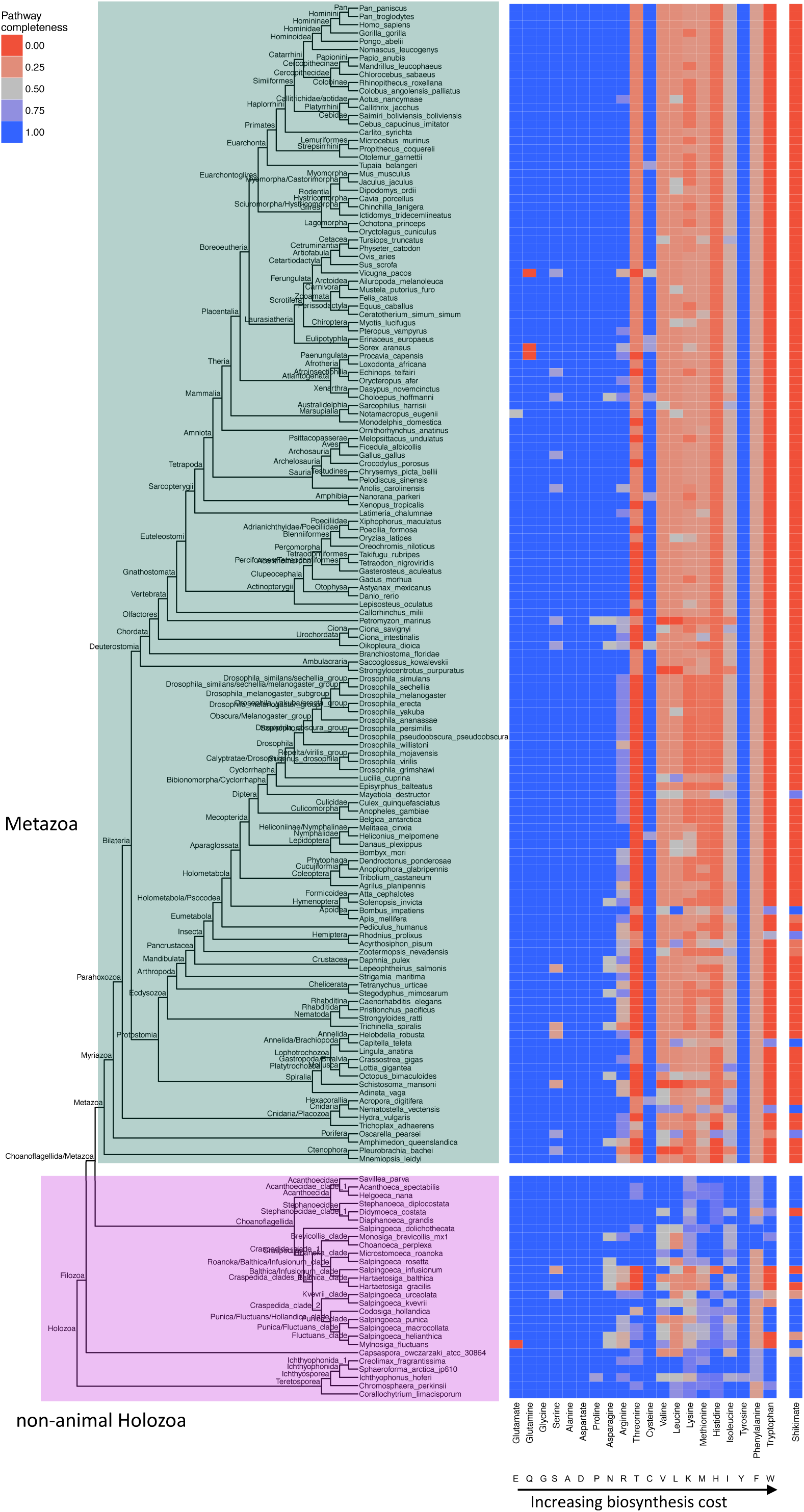
An overview of AA dispensability in Metazoa (fully resolved tree). We used the previously published database of 167 holozoans (Domazet-Lošo et al. 2024) to get a comprehensive overview of AA dispensability in this group. We retrieved all enzymes involved in AA biosynthesis from the KEGG and MetaCyc databases. We searched for their homologs within our reference database using DIAMOND (see Methods). For each AA, we showed the proportion of the enzymes with significant matches. In case of AAs with multiple alternative pathways, we showed the results only for the most complete one. Alongside the 20 proteinogenic AAs, shikimate pathway is also included as it is the precursor in the synthesis of aromatic AAs (histidine, phenylalanine, tryptophan, and tyrosine).

**Supplementary Fig. 6.**
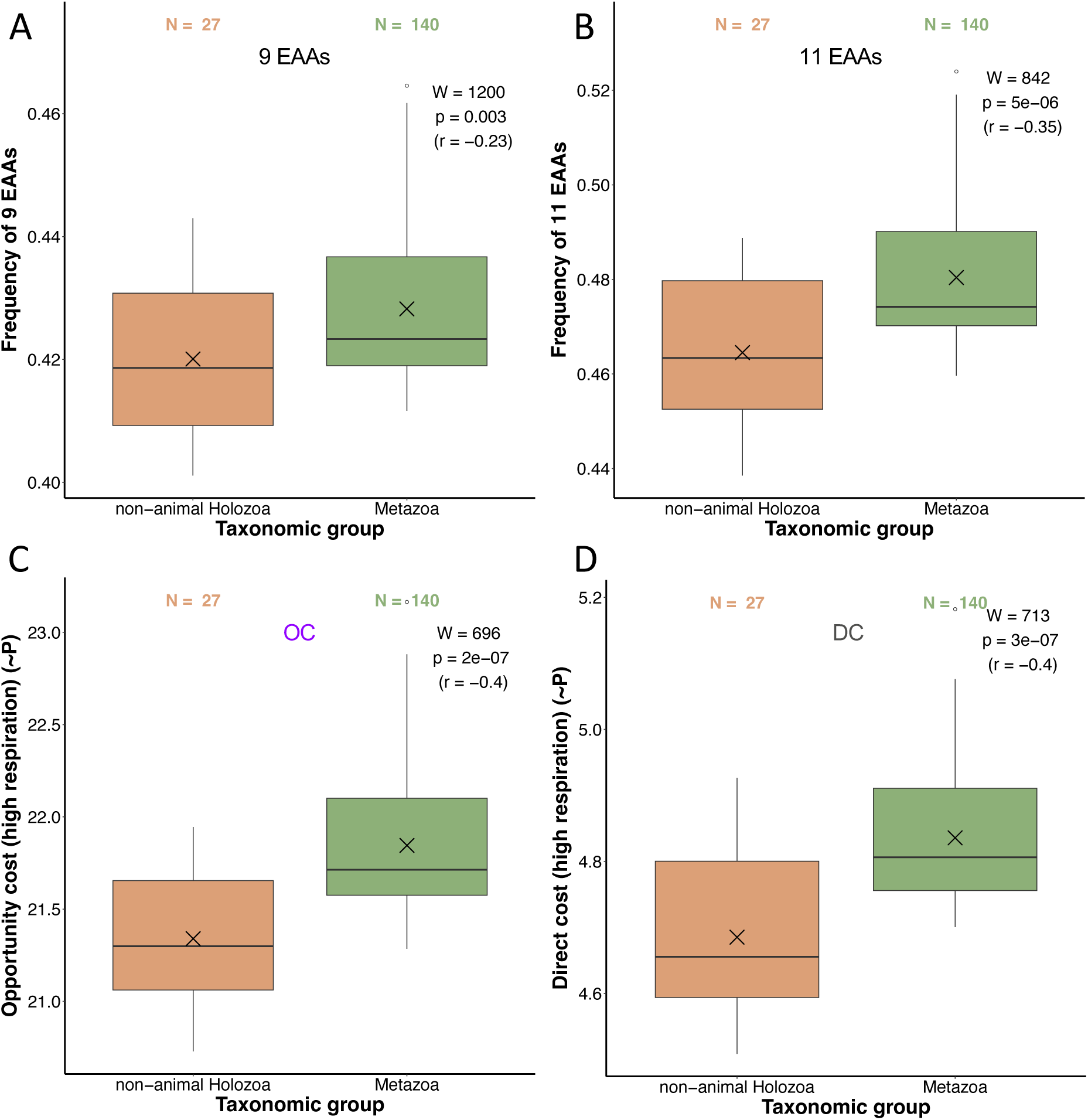
The comparison of EAA usage and the energy costs of an average AA in non-animal Holoza and Metazoa using only genes that are conserved between animals and non-animal holozoans. (**A**) The cumulative frequency of nine AAs which are generally considered to be essential in Metazoans (Thr, Val, Leu, Lys, Ile, Met, His, Phe, Trp). (**B**) The cumulative frequency of eleven EAAs. This extended dataset contains two AAs which could be considered conditionally essential (Cys, Tyr). (**C–D**) The opportunity cost (OC) and the direct cost (DC) of an average AA (see Supplementary Table 1). This value represents a weighted mean of AA biosynthesis energy costs where the frequencies of twenty AAs in a proteome act as weights. The differences in energy costs between the two groups were shown by boxplots and the significance of these differences was tested by the Mann-Whitney U test with continuity correction. We depicted the corresponding W-value, p-value, and effect size (r) in each panel. The X symbol represents the mean. The list of non-animal holozoans (27) and metazoans (140) whose proteomes are included in calculations is available in Supplementary Data 2 (Domazet-Lošo et al. 2024). To control for the presence of linage-specific genes, we first clustered all 167 proteomes using the following MMSeqs2 parameters: -e 0.001 -c 0.8 --max-seqs 400 --cluster-mode 1 --cov-mode 0 and obtained clusters of proteins with highly similar architectures. We then extracted only the clusters that contained both metazoan and non-metazoan holozoan proteins and then performed the frequency and energy cost analyses. The initial dataset contained 3,514,971 proteins, while the dataset with only conserved genes contained 1,460,551 proteins (42%).

